# The Human Octopus: Controlling supernumerary hands with the help of virtual reality

**DOI:** 10.1101/056812

**Authors:** Sander Kulu, Madis Vasser, Raul Vicente Zafra, Jaan Aru

**Affiliations:** Institute of Computer Science, University of Tartu, Tartu, Estonia

## Abstract

We investigated the “human octopus” phenomenon where subjects controlled virtual supernumerary hands through hand tracking technology and virtual reality. Four experiments were developed to study how subjects (n=10) operate with different number and behaviour of supernumerary hands. The behaviours involved inserting movement delays to the virtual hands and adjusting their movement scale or position. It was found that having more hands to operate with does not necessarily mean higher success rate while performinga certain task. However, supernumerary hands could be made more effective by adjusting the associated movement scales of the extra hands. The subjective feeling and ownership of the hands diminished when a delay was inserted for the virtual hands or when their position was altered.

## 1 INTRODUCTION

Humans have evolved from tetrapods, the first terrestrial animals with four limbs. The shape of the human body has transformed over time, but the number of limbs has remained constant. Humans have two lower and two upper limbs (i.e. legs and arms). But what if people had more than 2 arms? In these circumstances humans would potentially be able to perform more complex actions with less effort. In the present context we will call this phenomenon “the human octopus”.

Already from the ancient civilizations, people are known to believe in mythological creatures and gods that have multiple number of limbs. For example, the gods from Hinduism such as the governor of the Universe Vishnu,the warrior goddess Durga or the patron of arts and sciences Ganesha. The multiplicity of arms emphasized the deity’s immense power and ability to perform several acts at the same time. So the idea of having multiple limbs to operate with has lingered somewhere for thousands of years already. Modern computer technology is starting to provide us with tools that help to implement the idea of having multiple operable arms to some extent.

Recent advancements in computer science and stereoscopic displays allow us to create virtual experiences that can be perceived by humans as quasi-realistic. This approach opens up a whole range of new neurological and psychological experiments that have not been possible before. Furthermore, we can also monitor the movement of the body and limbs of the subject. This allows us for example to add two or more virtual arms to the virtual avatar that closely mimic the actions the subject does with his real hands. We could also deliberately insert a delay between the movement of real and virtual arms, adjust the hands movement scales and transforms.

This raises many interesting questions: If the subject had more than two arms to control, would he be able to perform actions more effectively inany way? How do humans solve the problem of controlling multiple pairs of arms with just one? To what extent would the subject perceive the virtual extra arms as his own and how would different manipulations affect that?

Several studies have shown that perceptual-motor and visuomotor synchrony(perceptual and visual stimulation that is in synchrony between virtual avatar and real body) is sufficient to create embodiment effects with virtual avatars [1] [2]. This means that participants tend to report the virtual bodies as the representation of themselves despite the fact that they know such experience is illusory. The subjects can also exhibit change in heart rate as a response to threats to their virtual avatar [3]. Psychological effects such as increased sweating have been detected upon the threats in virtual environment [4]. This means that virtual avatars are treated as a convincing representation of the subject’s own body in the virtual environment, which makes the virtual reality a viable tool for studying the perception of the human body schema.

Numerous studies have been conducted to learn about the perception of body in the brain through various illusions tapping onto body ownership [5]. The most known ones are the third hand and rubber hand illusions, but also invisible hand and out-of-body illusions. Experiments have shown that through the change of perspective it is possible to deceive the brain to accept the fake body parts as its own [6]. These studies reveal that reconstruction of the body in the brain can be somewhat manipulated under certain conditions such as the synchronous stimulation and the usage of arms that look very similar to our own. For example, in the third arm illusion, where another right arm is positioned next to the subject’s real right arm works only if the fake hand has the same color as the subject’s skin [4]. Virtual reality allows fulfilling most of the requirements that are necessary to deceive the brain about ownership of virtual limbs.

Several studies show that extended body avatars can be more useful than normal body avatars when they suit the task better. Firstly, there is research that investigated how subjects operate with 3-armed-avatar compared to 2-armed-avatar [7]. The third arm was designed to be significantly longer in order to reach further objects more easily. As expected, the subjects were more successful operating with 3-armed-avatar. Their following research that investigated embodiment effects of extended body avatars also claims that people can complete the task more successfully when the virtual avatar is not in one-to-one relationship with one’s real body [8]. Another interesting fact discovered was that participants performed more poorly with limbs that were not attached to the virtual body.

In addition to that, there is a study about controlling the supernumerary robotic hand by foot [9]. An experiment was designed to observe how subjects simultaneously catch three falling objects with both three and two hands setup. Once again it was found that three operable arms are preferred due to increased effectiveness. The reason behind it is that the task suits better for three hand setup rather than two hand setup. Furthermore, participants managed to control the third arm by foot without major issues.

There is another study that shows the strong relation between the individual’s vision and body perception. It claims that we do not perceive the environment itself but rather the relationships between our body and the environment [10]. Those relations determine the course of the most of our actions for instance grasping the objects or jumping over an obstacle. The article claims that if we would change the size of our hands or length of our feet, the brain would re-evaluate the relations between the object we are about to grasp and our increased size hands [10]. This means that our brain is able to adapt to the changes and perceive the size of our increased arms correctly. In the current study it is presumed that similaradaptions may occur when a delay is inserted into the hands or their movement scale is changed.

The present work focused on developing a set of experiments involving different hand setups that were used to complete a specific task over a short period of time. Four experiments were conducted on a small sample in order to test the applicability and get initial data about the above questions. These four experiments are part of a Unity Engine project which is freely available to assist anyone in conducting further studies.

## 2 MATERIALS AND METHODS

### Sample

In total 10 subjects, 8 males and 2 females, volunteered to take part in the experiments. Eight participants were right handed and two participants left handed. Their log files and questionnaires were used to examine each of the four experiments separately. The results are supported by observational findings and participants’ comments.

### Hardware

We used the Oculus rift DK2 virtual reality headset with low-latency positional tracking, OLED screens (960×1080 pixels per eye resolution) and 110 degree field of view[11] [12]. Hands of the subjects were tracked using the Leap Motion hand tracker peripheral. It uses two monochromatic cameras and three infrared LEDs to observe the hemispheric area up to1 meter [14]. The hardware in combination with dedicated software provides robust and low latency hand recognition [15]. The program ran on a pc with a i7 3.5GHz CPU, GeForce 980 Ti GPU and 16 GB RAM.

### Software

Unity game engine was used for developing the software [17]. Unity has both virtual reality and Leap Motion compatibility which made it a suitable candidate to create our experiments with.

### Leap Motion code modifications

Leap Motion’s Unity core assets package is mainly meant to display only one pair of hands, so various modifications were done to the existing assets. Each HandController component was made to represent only one hand, enabling to change the parametrical values of each hand separately. For logging purposes, a name variable was also added to a HandController class.

For each frame in Unity scene HandController class receives a hand image that represents the current position of the real hands. In order to add the delay into the hands, a hand image buffer was created and added to the HandController instance. Depending on the desired delay certain amount of hand images are buffered into a list in order to use them after the right amount of frames has passed.In order to bind head movement to the hands, one simply has to position the HandController more forward from the camera. When Leap Motion is attached to the head mounted display, its software compensates the head movement by simply moving the hands in opposite direction. If HandController is positioned right in front of the camera in the scene, the amount of compensation is correct and one’s hands seem to stay in one place when the head moves. But when HandController is positioned further, the compensation becomes too great which makes the hands move with one’s line of sight.

HandController class has equivalent editor script to enable changing values from Unity’s inspector/editor. By default, it has variables for position, size and movement scale. Option to choose amount of frames fora delay was added.

### Virtual Environment

In order to create the experiment itself, a script was written to manage the falling object. Attaching it to an object makes it fall from certain height at random position. For this experiment a small green cube was created. The object is positioned to a rectangular area which can be adjusted to the liking. If the object reaches certain height or collides with any of the hands it respawns at newly randomized position. At each frame a velocity vector is given to the object to make it fall at constant speed.

Each experiment can be divided into phases that have different velocity of falling. There are also pauses between phases because keeping your real hands in Leap Motion’s tracking space for more than a minute can be tiring. Experiment starts when the space key is pressed and ends when the time is up.

The script is also responsible for playing sound effects when the object collides with the hand or when the experiment starts or ends. Sound effects are taken from https://www.freesound.org and are licensed to use pretty much in any way that users want as long as the source is referenced [18][19][20].

The environment of the scene is a room that has minimalistic furnishingof various classroom prefabs, as can be seen on figure 1. These prefabs are available for free from Unity’s assets store [21]. Purpose of such interior is to make the environment feel more natural but not too overwhelming for the subjects. Virtual hands and the camera are positioned front of the table in the middle of the room. Purpose of the table is to keep the subjects from holding their hands too low in order to accomplish better tracking. A wooden box is positionedabove the table and it marks the area in which the cube is respawned.

**Figure 1:**
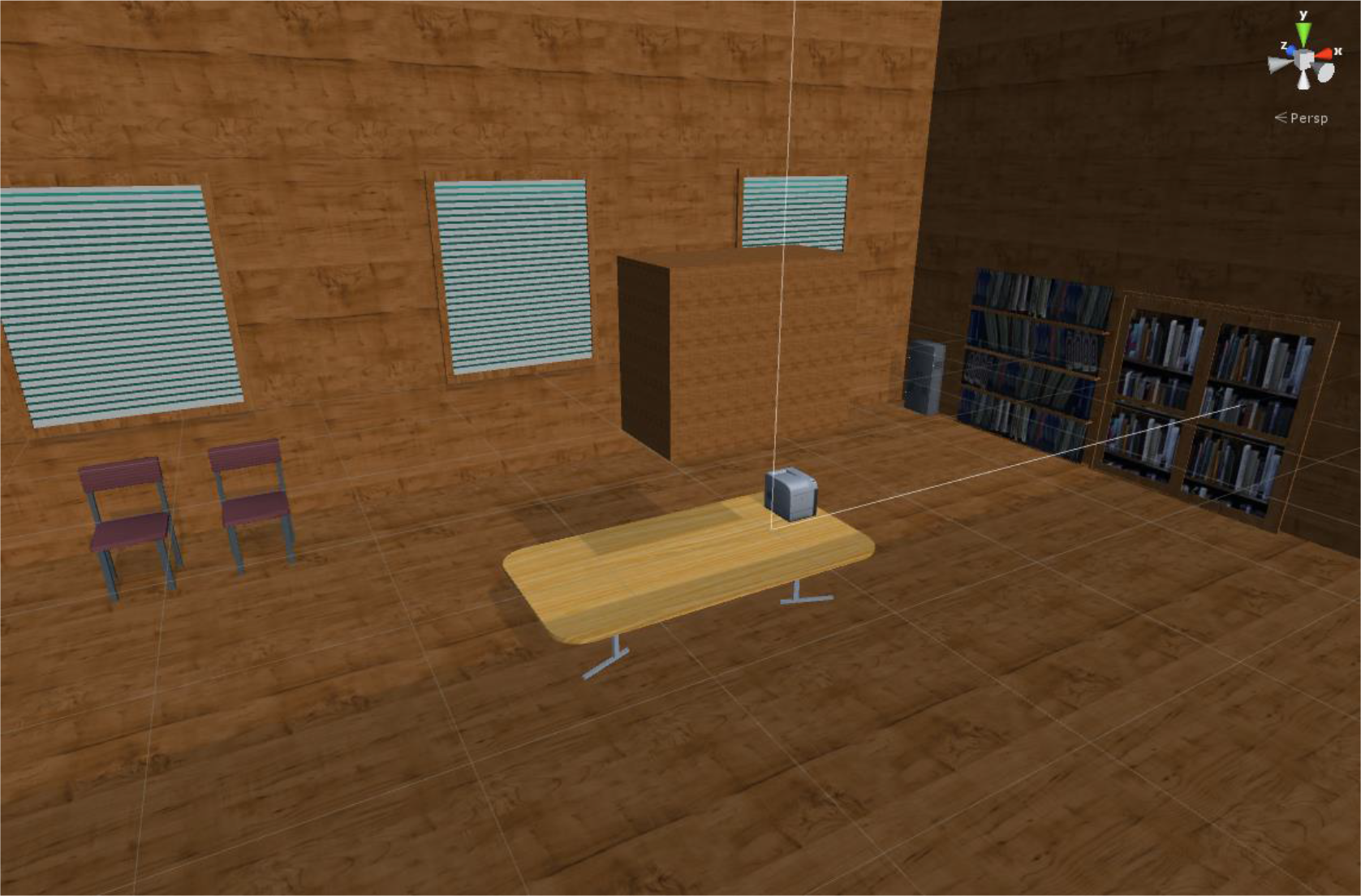
Environment of the scene

### Experimental task and procedure

The task was to catch the object (a green cube) that is dropped from a box above the subject’s virtual avatar. The box marks the rectangular area of possible drop positions for the object and can be seen when the subject looks up. The object is returned to the box after the subject has managed to touch it with any of his hands or when the object hits the ground. The position of the object is randomized each drop. Distance between the spawn height and reset height was 2 Unity units, corresponding to approximately 2 meters in real word scale. Subject’s view of the scene can be seen on figure 2.

**Figure 2:**
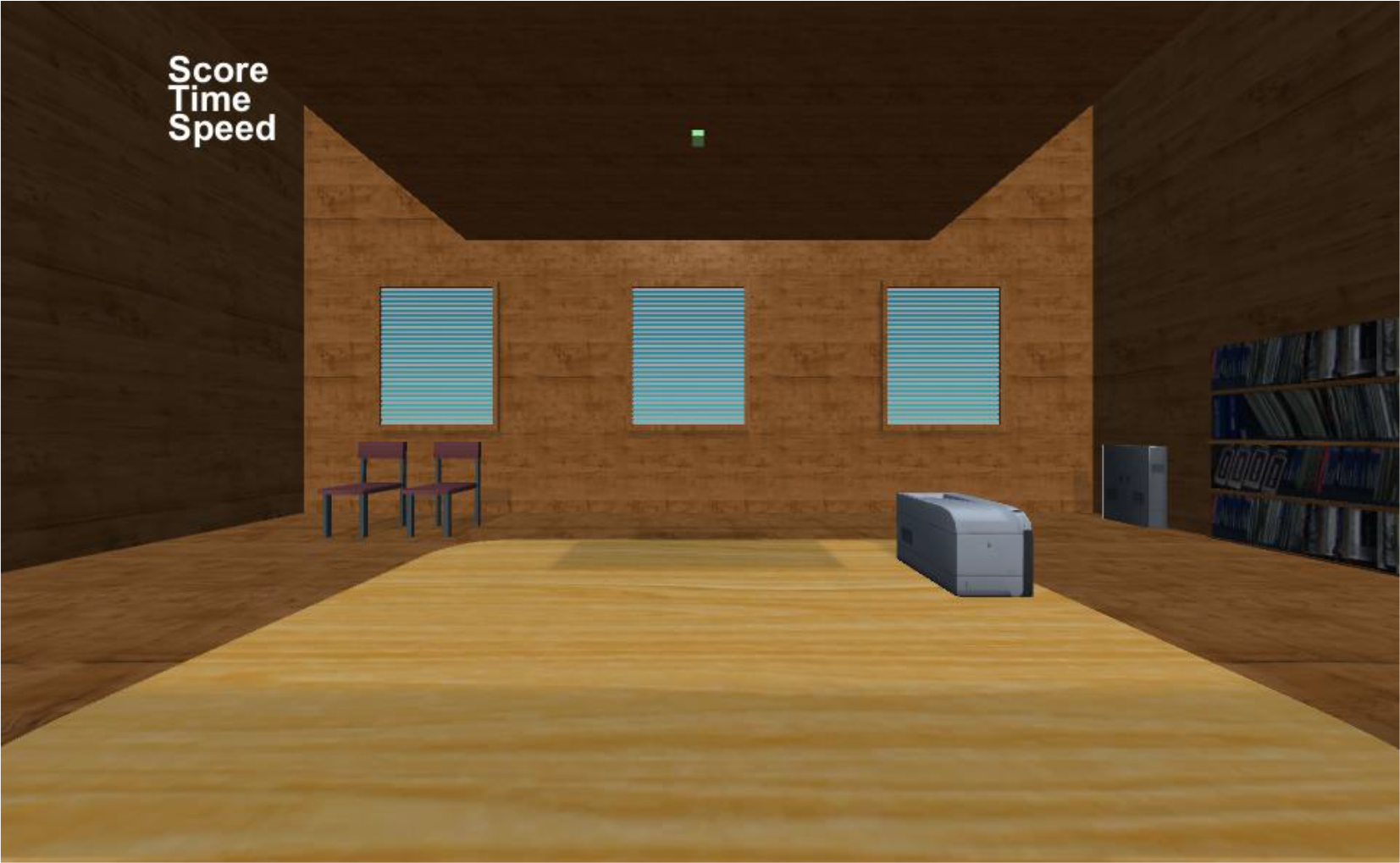
Subject’s view without the hand avatars.

The task was divided into three phases, each lasting 60 seconds. There was a 30 second resting pause between the phases. The falling speed of the green cube had progressive difficulty, determined by a pilot study. In the first phase the cube dropped 0.5 units per second. In the second and third phase the speed was significantly higher, 1.3 and 1.9 units per second respectively. This made the task much more difficult and required fast and precise movement of hands.

The subject received auditory feedback when a catch was successful, andin the beginning and end of every phase.

The task was accompanied by questionnaires about different aspects of the hand setups, filled after each experiment. Each questionnaire consisted of grading the statements given below using a Likert type scale ranging from 1 to 5, where 1 stands for strong disagreement, 3 for neither agreement nor disagreement and 5 for strong agreement. In addition participants had an option to give verbal feedback and other comments about the experiments.For the full questionnaire see appendix 1.

Experiments were administered in the same order as described below and each participant completed each experiment.

### Different hand setups

For the current experiments the hands only model was preferred over full length arms. This was to prevent subjects randomly hitting objects with their forearm or elbow. The hands had given humanlike graphics model which had the similar skin tone of the subject.

### First experiment

In the first experiment subjects had one pair of virtual arms (figure 3) that moved almost exactly as one’s real arms except that the virtual arms movement scale was slightlyincreased (1.4 times). This allowed to increase the reach for further awat objects. Otherwise the task would have been impossible compared to otherhand setups.

**Figure 3:**
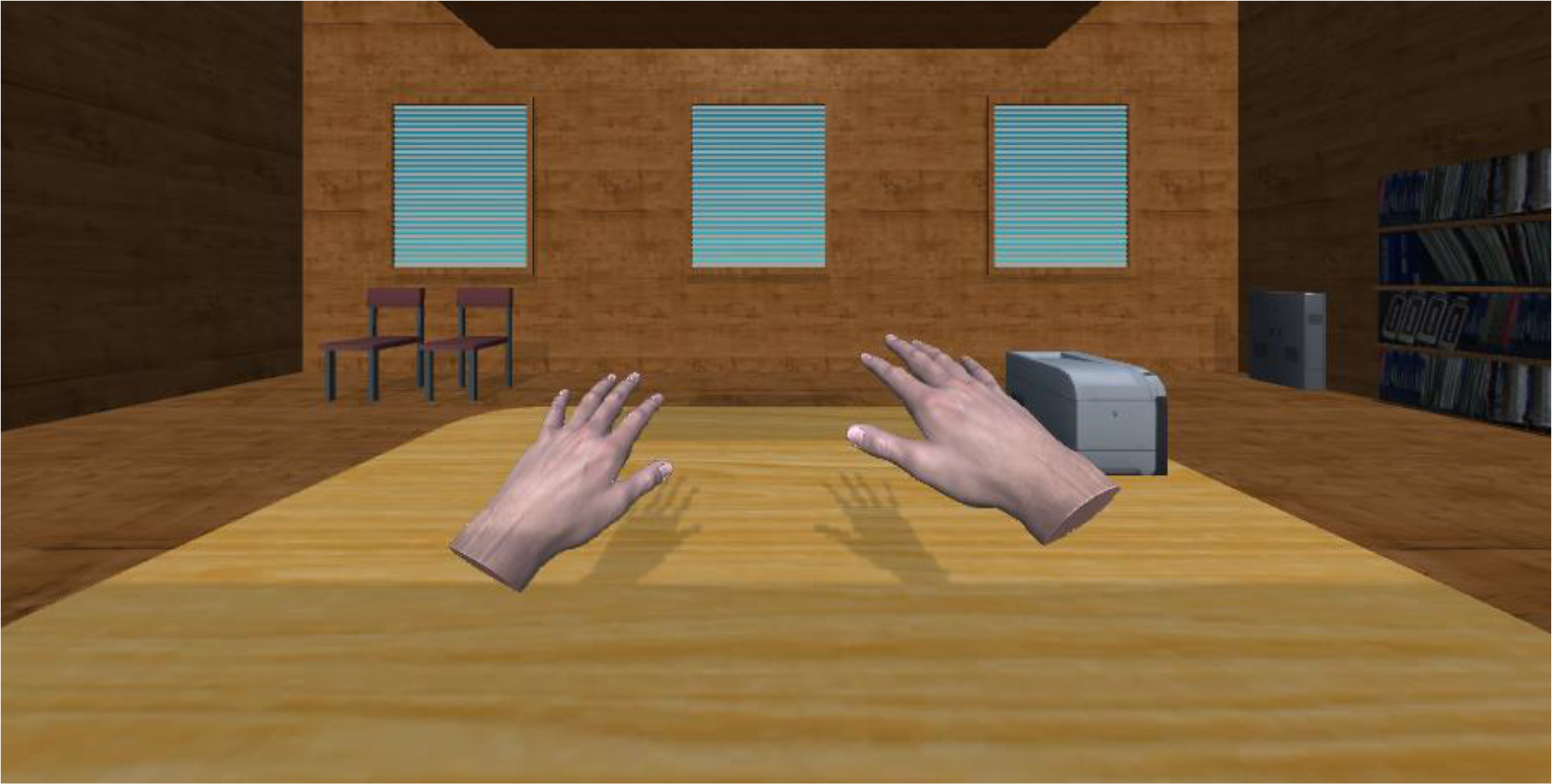
Hand setup of the first experiment.

### Second experiment

The second experiment examines how the subject completes the same task with increased number of hands. Subject is given three pairs of arms (figure 4) which all move simultaneouslyto one’s real arms. All arms are lined up horizontally and each pair has a different value of movement scale depending on the position – inner pair of hands has normal (1.0), middle one has slightly increased (1.5) and outer pair significantly increased (2.0) movement scale. This means that each pair has a different field of grasp depending on its distance from themiddle. Subject’s task is to catch falling objects with any of one’s arms.

**Figure 4:**
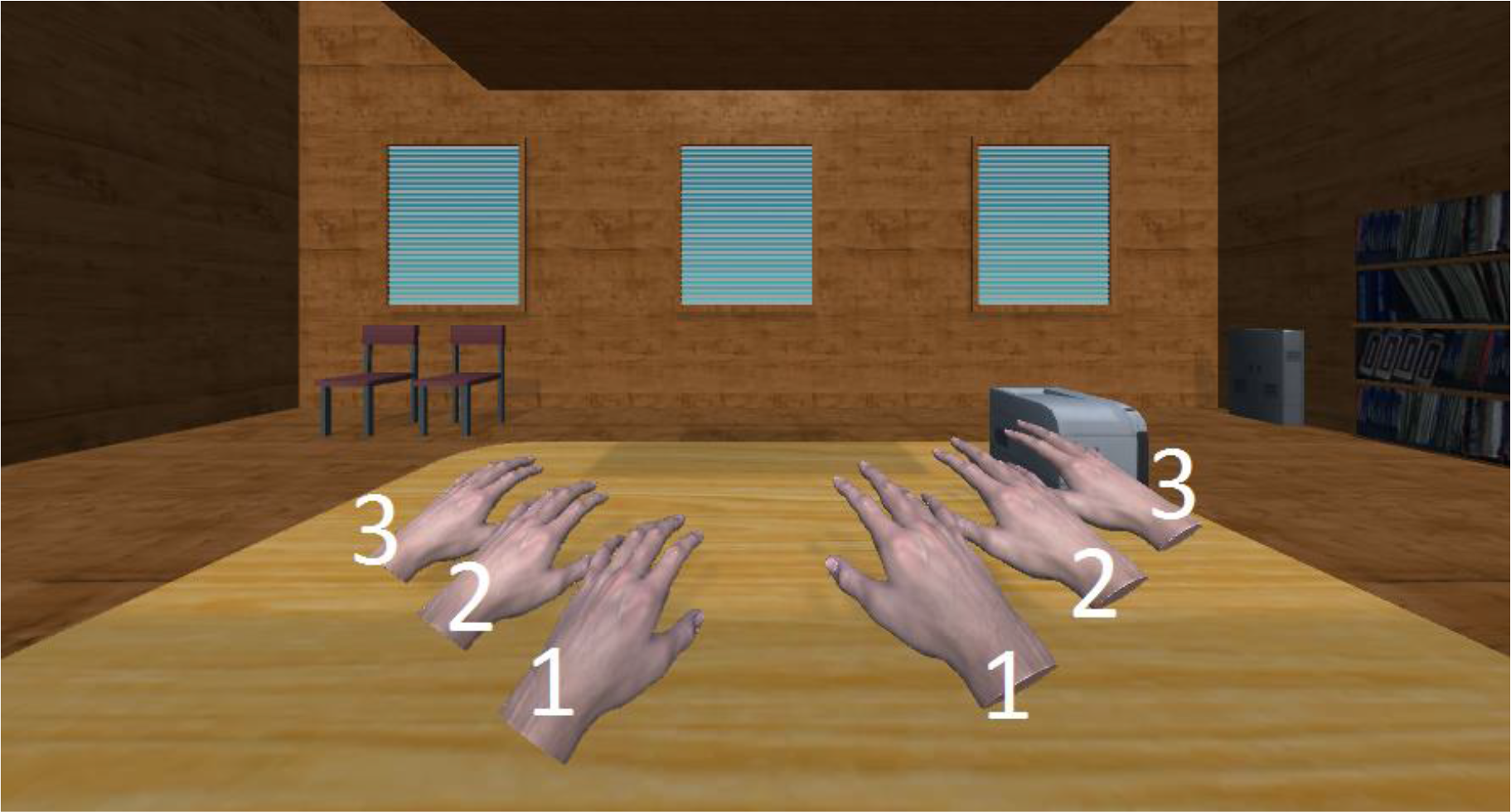
Hand setup of the second experiment. Numbers mark the number of pair.

### Third experiment

The third experiment has the same number and movement scales of arms assecond experiment, but there is a slight delay in the hand movement depending on the position. The inner pair of arms moves simultaneously to the one’s real arms and is considered to be the main pair. Two other pairs have a slight delay depending on the distance from the middle. Second pair has a 15 frames delay and third pair has a 30 frames delay. The task remains the same but now the subject has to adapt to the totally different hand movement. Hand setup of the third experiment can be seen on figure 5.

**Figure 5:**
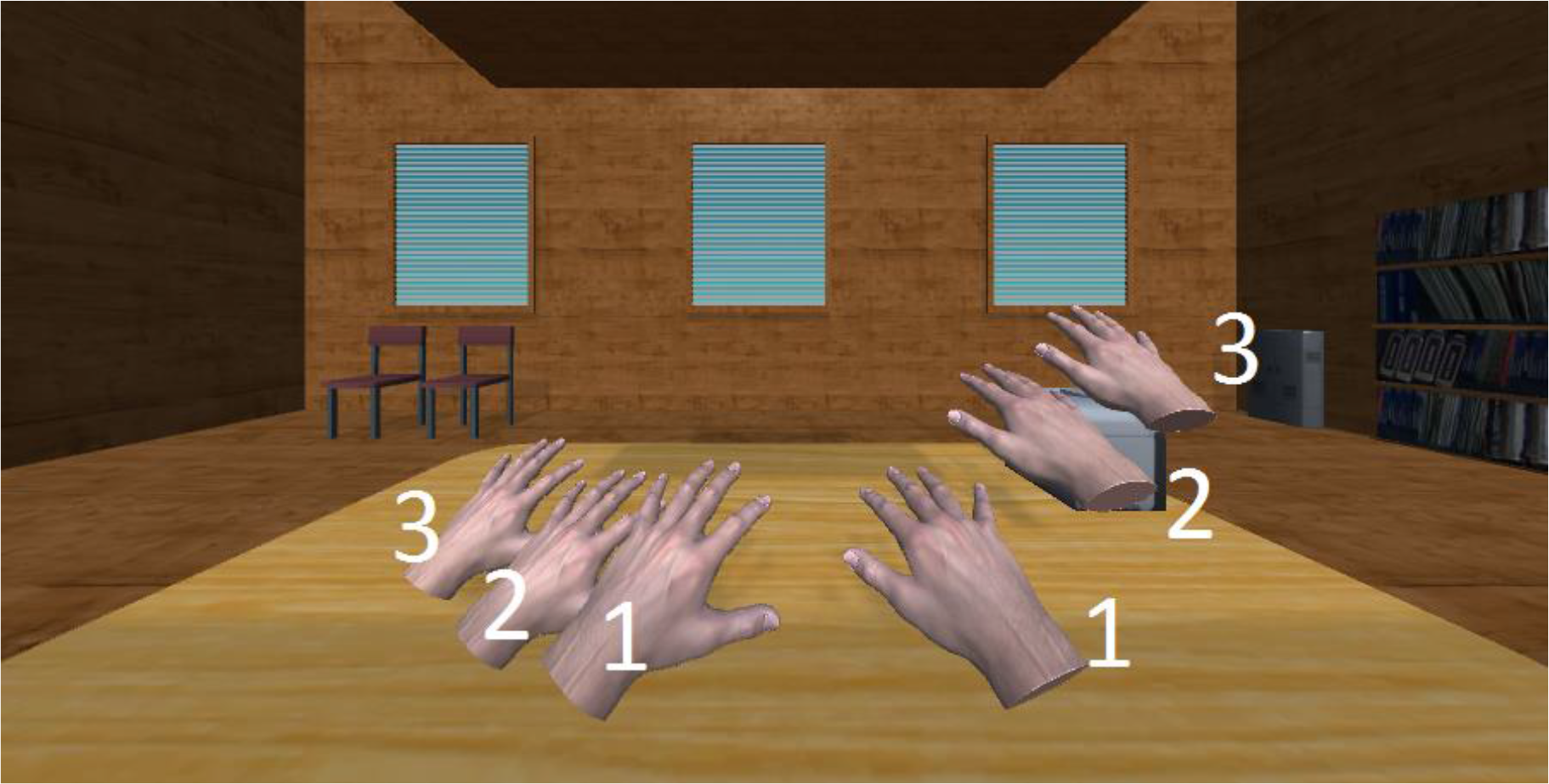
Hand setup of the third experiment. Numbers mark the number ofpair.

### Fourth experiment

The last experiment examines how the subject adapts to the control strategy that involves both hand and head movement. Subject has total of four pair of hands that are all in different position and have different rotation as can be seen on figure 6. First pair of arms is located and rotated as one’s real arms and is considered to be the main pair. Second pair of arms is located slightly forward to the left and rotated to face toward the center of area in which the objects are falling. Third pair of arms is similar to the second pair except it is located on the right. The last pair is located horizontally in the center like first pair but is significantly further forward so that it is on the other side of the area in which the objects are falling. Last pair is also rotated to face the center of the area so to a subject it looks mirrored. All pairs of arms except the first are bound to subject’s line of sight. This means that subject can move one’s virtual arms with head movement. If subject turns one’s head left or right the arms move left or right respectively as well. Arms are also bound to move similarly when subject tilts the head left or right or when subject moves head forward or backwards. The further arms are the greater is the movement created by head tracking.

**Figure 6:**
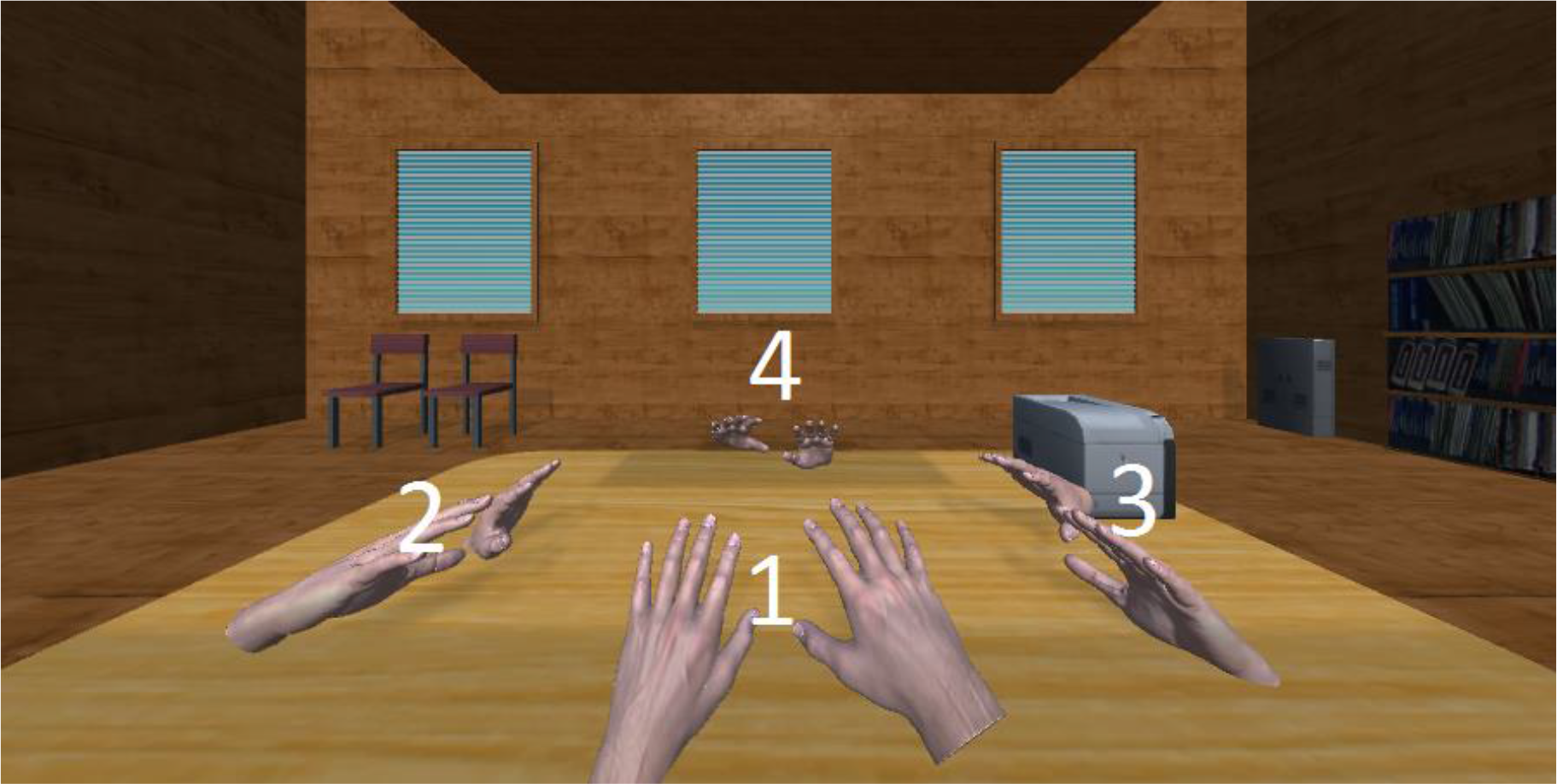
Hand setup of the fourth experiment. Numbers mark the number of pair.

### Experimental procedure

Subjects were asked to take part in a virtual reality experiment which involves operating with various supernumerary hands. Each subject is told/shown the short description of the task which can be found below. After reading the short description the subject is asked to sit on a chair. After that it is made sure that subject has enough space to move one’s hands freely. Subjects also receive thorough instructions to accomplish besttracking possible 3 with Leap Motion. If subject is ready he is asked to put on the Oculus Rift and headphones. The experiment begins with training phase where subject can exercise with current hand setup. When the subject understands the setup and is ready to proceed, the task is started. After completing the task, the subject is asked to assess a few statements (see below) towards body perception based on a Likert-type response scale. If the subject is ready to proceed, he or she is asked to put on the Oculus Rift again and the next experiment is started. In this manner all the experiments will be done with short breaks between them.

## 3 RESULTS

### 3.1 First experiment

In the first experiment where subjects had only one pair of hands, subjects were relatively successful at catching falling objects. In the first phase 70% of the subjects managed to catch all the cubes. The mean score for the first phase is 96% with standard deviation (SD) of 7.1, which means that subjects were almost equally successful.

In the second phase results between subjects differ the most with mean score of 64% and SD of 17.5. As expected the results dropped even morein thethird phase where the mean score is 31.3% with standard deviation of 15.6.

When asked if the hands felt as if they were one’s own hands, subjects mostly agreed or were neutral. No certain strategies were developed except the usage of hands as one would normally do. Some subjects also realized during the experiment that holding hands higher grants better tracking despite the fact that it was told before the experiment.

### 3.2 Second experiment

In the second experiment subjects had three pairs of hands with different movement scale and the results are actually very similar to the first experiment with an exception of the last phase. Score in the first and second phase are 96.9±3% and 64.64±14.4% respectively. In the last phase subjects managed to catch almost 10% more objects (40.9±15.9%) compared to the first experiment (31.3±15.6). This might be due to a fact that subjects had really hard time catching objects at such high speed and having an increased number of hands simply increased the area that subjects can cover under the drop zone.

Since each pair of hands had different movement scale it is worth observing which hands were used the most. It appears that at higher speeds subjects mostly tended to catch objects with the second pair of hands which had asimilar movement scale as hands in the first experiment. Surprisingly in the last phase the right hand is preferred much more compared to the first experiment.

Although the success rate is very similar to the first experiment all subjects agreed that extra hands were helpful completing the task (4.7±0.5%). Observational findings revealed that subjects moved their hands much less since each pair of hands reached a different distance.

Subjects also agreed that all three pairs of hands felt equally as their own hands (4.2±0.6%). Although the whole experiment took about 5 minutes, most of the subjects claimed that they became better in time controlling the extra pairs of hands (3.8±1.5%).

When asked about strategies that subjects developed most of them claimed to have no certain strategy. Nevertheless, a few strategies were still developed such as juggling your hands back and forth really fast or using mainly outer pair of hands due to its higher movement scale.

### 3.3 Third experiment

In the third experiment subjects had three pairs of hands with different movement scale and delay. Success rate seems to decrease when a delay is inserted into the hands. In the first and second phase the success rates are 89.6±8.4% and 59.1±13.2% respectively which is about 6-7% lower when compared to the first and second experiment. Notably in last phase, subjects still showed better results (38.9±9.6%) than in the first experiment (31.3±15.6%). It is similar to the second experiment’s last phase.

For the most of the subjects the difference between first and second phase is greater than between second and third, which is not the case in second experiment. This shows that delayed hands tend to be useful only when objects are falling slowly. It makes sense because at higher speeds the time window of catching the object gets closer to the amount of delay that is inserted in to the hand.

Observational findings reveal that subjects mostly tried to use their main pair of hands that did not have a delay. That is supported by subjects’ comments that mostly claimed that having a delay in hands is a confusing factor and therefore they mostly preferred using the main pair of hands that did not have a delay. However, in the last phase subjects actually mostly used the second pair.

Although most of the subjects disagreed (2±1.4) about strategy development, a few strategies were still developed with delayed hands. Firstly, two subjects claimed that keeping their hands far forward would keep delayed hands in better position to catch further objects. Therefore, bigger movements were required only to catch closer objects with hands that did not have delay. Second strategy is about using delayed hands like a whip due to a fact that delayed hands would follow main pair of hands shortly after. Such strategy requires predicting the future hand position and indicates that delayed hands might be treated as tools rather than own hands.
It is supported by the fact that subjects mostly disagreed (2±0.8) about feeling that all hands are equally their own and were neutral (3±1.2) about the feeling that delayed hands became more natural in time.One subject claimed that delayed hands rather felt like an extension of main pair of hands and used them as an extra tool.

### 3.4 Fourth experiment

In the last experiment subjects had four pairs of hands in different positions and success rate is the worst in all phases having the scores of 78.89±12.4%, 51±9.2% and 31.1±11.3% respectively for three phases. However, in the last phase the success rate is almost equal to the first experiment (31.3±15.6%).

Hands on the left and right were designed to catch objects that are falling from the side but most of the subjects found that they were mostly useless. Second and third pair were used least. This indicates that hands onthe side could have been positioned better when designing the experiment.

Fourth pair of hands was mirrored towards subject’s view therefore making it more confusing to control. Despite that fourth pair was still used to catch most of the objects with an exception of first phase as can be seen on hands used graph. Reason behind that might be the factor that one could use head movement to assist controlling the fourth pair.

Observational findings reveal that almost all subjects used their head movement to assist controlling the supernumerary hands at one point. When asked if head movement was preferred over hand movement the results were different for most of the subjects, but the average (2.7±1.1) shows that general opinion was almost neutral. Similar result (2.5±1.4) was detected for natural feeling when controlling arms with head movement.

Subjects disagreed (1.9±0.7) about feeling that hands in other position felt as if they were one’s own hands. It was also claimed that such hand setup is too confusing and is almost impossible to make fast and precise movements in order to catch objects.

As can be seen by relatively low standard deviations, subjects are not as much scattered than in the previous experiments. This might also indicate that such hand setup is really confusing, which makes the skill differences of controlling supernumerary hands diminish.

General opinion about developing certain strategies is almost neutral (2.6±1.7). However, a few strategies were still described. The most popular strategy was to cluster all four pairs of hands together and then control only with head movement. However, it was also claimed and observed that such strategy did not work too well. Second strategy was to just use main pair of hands as much as possible.

### 3.5 General results

When inspecting averages of all the experiments on one graph (figure 7)it can be seen that subjects performed best in the second experiment, where they had three pairs of hands with different movement scale and without a delay.

If we compare the setups with supernumerary hands to just two hand setup we can see that success rate drops less with increased number of hands as the speed increases. This is probably due to the fact that subjects have higher chance to catch an object with increased number of hands than with just one pair.

It was also claimed that in the second experiment supernumerary hands were helpful completing the task. Furthermore, when stated that having more arms to operate with helps to complete the task more successfully over all the experiments with supernumerary hands, the general opinion slightly leans toward agreement (3.3±1.2). Similar result (3.1±1) was detected when asked if subjects became better in time when controlling extrapair of hands in general for all experiments with supernumerary hands. Although almost all subjects claimed that it was different for each hand setup.

**Figure 7:**
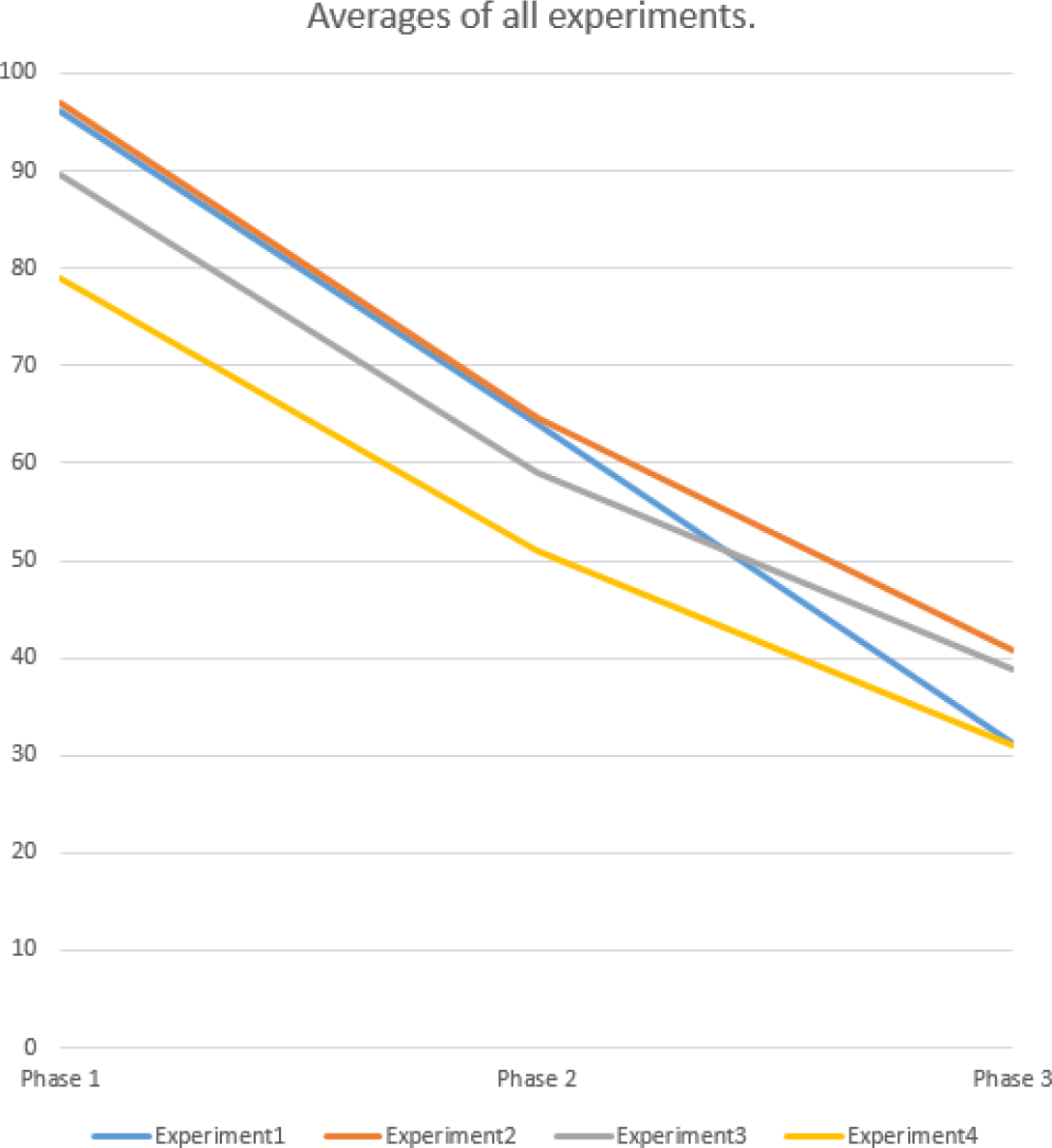
Averages of all experiments. Y-axis shows the average percentage of caught objects.

In the last experiment the performance is much poorer than in other experiments despite having the highest amount of arms. Controlling four pairs of hands that are all in different position requires a lot of coordination. Although supernumerary arms had head movement to assist controlling them, most participants still preferred to move them naturally with real hands. Some claimed that having such hand setup is too confusing specially when it requires fast reactions at higher speeds.

## 4 DISCUSSION

Using virtual reality, we investigated the “human octopus” phenomenon. It was found that having more arms to operate with does not necessarily mean that one would perform the task more successfully. However, the subjects had relatively small amount of time to practice. It was noticed during the development of the experiments that if one operates with certain hand setup for longer period of time, one gets significantly better at controlling increased number of hands.

Although adjusting the movement scale of hands could make them more effective resulting in less hand movement, it is highly dependent on the task. It is worth observing how subjects would perform on more than just one task. Altering the behaviour of hands could really improve performance at some specific tasks.

The natural feeling and ownership of the hands seems to diminish when adelay is inserted into them or their position is altered. It was stated that such manipulation gives the feeling that supernumerary hands are rather an extensive tool than part of their body schema. Although such manipulation made controlling supernumerary hands rather clumsy, they were still somewhat helpful towards completing the task.

Another interesting aspect was detected as the subject (S7) who claimed to play a lot of video games that require fast reactions, has the best result in almost every phase throughout all four experiments even though it was his first virtual reality experience. Reason behind that might be that people who play competitive video games tend to have better reaction time and easier time adapting to new virtual environments.

It is also worth mentioning, that even though the supernumerary hands were not most effective, almost all of the subjects found the experience to be ‘really trippy’, interesting, cool and entertaining overall.

When it comes to involving head movement to control strategy, it is definitely worth researching more into. Although in current experiment, subjects were almost neutral about preferring head movement over hand movement,almost all the subjects used head movement to assist controlling supernumerary hands at one point. However, the question about effectiveness of head movement based control strategy remains unanswered.

Observational findings and opinions of participants were also used to evaluate applicability of the experiments. Several shortages were discovered when carrying out the experiments.

Firstly, it is really difficult to find a speed that suits perfectly for all subjects. In the current experiments three different velocities were used. In the first phase the speed was slow enough for subjects to catch about 90% of the objects. In second phase, which seemed to suit the subjects most, the success rate was around 60%. However, in the last phase least successful subjects had really tough time catching the objects, reaching the success rate around 10%. Therefore, for further research it is suggested to increase the number of phases and velocities to suit the intraindividual variability of the subjects better. Another approach would be that speed of the object increases during the experiment when subject manages to catch it. In this manner every subject could develop the speed that suits him or her best.

Secondly, there were some problems with hand tracking. All subjects encountered the disappearance of hands at some point leading to increased number of missed objects. It was also observed that even though each subject received the best instructions possible about tracking, at higher speeds subjects tended to move their body too eagerly or keep their hands beneath the hemisphere of Leap Motion therefore causing the loss of tracking. During the process of development, it was found that it takes serious amount of experience to truly understand the tracking capabilities of Leap Motion.
Furthermore, in current experiments the area in which the objects dropped was rectangular. This caused a few occasions where subjects caught only glimpse of the object or did not see it at all because they were looking on the other side or leaned too much forward. Therefore, another shape of area is recommended to use, such as circular or sectorial shape. Ideally the size of the area should also be in relation to subjects’ hand length.

It was also stated by subjects and confirmed by observational findings that the task in current experiments requires fast reactions. This might draw focus of this experiment away from observing how subjects operate with supernumerary hands to observing how fast the subjects react.

It is strongly recommended to address these problems before applying them on larger scale or conducting further researches. In conclusion, it is difficult to design the task in a way that suits all the subjects.

In conclusion it can be said that having more arms to operate with doesnot necessarily mean that one would perform more successfully, especially when their control strategies are still reliant only on one pair of real hands. However supernumerary hands can still be helpful even when they all move simultaneously to real hands resulting in less total hand movement. In addition to that, increasing the movement scale of hand, enabling it to reach further, makes it more effective when it suits the task better. Even if the idea of owning supernumerary hands will not turn out to be effective in a practical way in future, it has great potential in the entertainment industry.

## Appendix 1 Questions in the questionnaires

Experiment 1: 
- Controlling one pair of hands felt as if they were my own hands.

Experiment 2: 
- Controlling three pairs of hands felt as if they were all equally my own hands.
- Extra pairs of hands were helpful completing the task.
- I became better in time at controlling extra pairs of hands.
- I developed a certain strategy to complete the task using the supernumerary hands. If yes, explain it.

Experiment 3: 
- Controlling three pairs of hands with a delay felt as if they were all equally my own hands.
- The feeling of having delayed hands became more natural in time.
- I learned to use delayed hands in a useful way.
- I developed a certain strategy to complete the task using the supernumerary hands. If yes, explain it

Experiment 4: 
- The hands in other position felt as if they were my own hands.
- It felt natural to control arms with head movement.
- I rather controlled virtual arms with head than real hands.
- I developed a certain strategy to complete the task using the supernumerary hands. If yes, explain it.

After all experiments are completed: 
- Having more arms to operate with helps to complete the task more successfully.
- Controlling supernumerary hands felt as if they were all equally my own hands.
- I became better in time when controlling extra pairs of hands.
- I felt physical burden when controlling supernumerary hands.
- I felt mental burden when controlling supernumerary hands.

## REFERENCES

[1] Sanchez-Vives, M. V., & Slater, M. (2005). From presence to consciousness through virtual reality. Nature Reviews Neuroscience, 6 (4), 332–339.

[2] Sanchez-Vives, M. V., Spanlang, B., Frisoli, A., Bergamasco, M., & Slater, M. (2010). Virtual hand illusion induced by visuomotor correlations. PloS one, 5 (4), e10381.

[3] Slater, M., Spanlang, B., Sanchez-Vives, M. V., & Blanke, O. (2010). First person experience of body transfer in virtual reality. PloS one, 5 (5), e10564.

[4] Guterstam, A., Petkova, V. I., & Ehrsson, H. H. (2011). The illusion of owning a third arm. PloS one, 6 (2), e17208.

[5] Guterstam, A., Gentile, G., & Ehrsson, H. H. (2013). The invisible hand illusion: multisensory integration leads to the embodiment of a discrete volume of empty space. Journal of cognitive neuroscience, 25 (7), 1078–1099.

[6] Ehrsson, H. H., (2004). Experiments with a rubber hand reveals how the brain recognizes it’s own body. J Swedish Med. Assoc., 48: 3872.

[7] Won, A. S., Bailenson, J., Lee, J., & Lanier, J. (2015). Homuncular flexibilityin virtual reality. Journal of Computer-Mediated Communication, 20 (3), 241–259.

[8] (2016, May) Won, A. S., Bailenson, J. N., & Lanier, J. Appearance and Task Success in Novel Avatars. Visited 11.05.2016 [Online] Available: https://vhil.stanford.edu/mm/2015/12/won-presence-task-success.pdf

[9] Abdi, E., Burdet, E., Bouri, M., & Bleuler, H. (2015). Control of a Supernumerary Robotic Hand by Foot: An Experimental Study in Virtual Reality. PloS one, 10 (7), e0134501.

[10] Linkenauger, S. A., Leyrer, M., Bulthoff, H. H., & Mohler, B. J. (2013). Welcome to wonderland: The influence of the size and shape of a virtual hand on the perceived size and shape of virtual objects. PloS one, 8 (7), e68594.

[11] (2016, April) Oculus rift. Oculus Rift. Visited 20.04.2016. [Online]. Available: https://www.oculus.com/en-us/dk2/

[12] (2016, April) Oculus Rift.Wikipedia. Visited 20.04.2016. [Online]. Available: https://en.wikipedia.org/wiki/OculusRift

[13] (2016, May) Oculus Rift Development Kit 2. Wikimedia Commons. Visited 01.05.2016. [Online] Available: https://commons.wikimedia.org/w/index.php?curid=35919898

[14] (2016, May) How Does the Leap Motion Controller Work? Leap Motion. Visited 09.05.2016. [Online]. Available: http://blog.leapmotion.com/hardware-to-software-how-does-the-leap-motion-controller-work/

[15] (2016, April) Orion Beta. Leap Motion. Visited 21.04.2016. [Online]. Available: https://developer.leapmotion.com/orion

[16] (2016, May) Leap Motion Controller. Wikimedia Commons. [Online]. Available: https://commons.wikimedia.org/w/index.php?curid=48288498

[17] (2016, April) Unity.Unity Technologies. Visited 21.04.2016. [Online]. Available: https://unity3d.com/unity

[18] (2016, May) Countdown Sound. Freesound. Visited 10.05.2016. [Online]. Available: https://www.freesound.org/people/meatball4u/sounds/17216/

[19] (2016, May) Bell Sound. Freesound. Visited 10.05.2016. [Online].Available: https://www.freesound.org/people/CJ4096/sounds/66717/

[20] (2016, May) Game Sound Correct. Freesound. Visited 10.05.2016. [Online]. Available: https://www.freesound.org/people/Bertrof/sounds/131660/

[21] (2016, April) Props for the Classroom. VR. Visited 24.04.2016. [Online]. Available: https://www.assetstore.unity3d.com/en/#!/content/5977

